# Suboptimal phenotypic reliability impedes reproducible human neuroscience

**DOI:** 10.1101/2022.07.22.501193

**Authors:** Aki Nikolaidis, Andrew A. Chen, Xiaoning He, Russell Shinohara, Joshua Vogelstein, Michael Milham, Haochang Shou

## Abstract

Biomarkers of behavior and psychiatric illness for cognitive and clinical neuroscience remain out of reach^1–4^. Suboptimal reliability of biological measurements, such as functional magnetic resonance imaging (fMRI), is increasingly cited as a primary culprit for discouragingly large sample size requirements and poor reproducibility of brain-based biomarker discovery^1,5–7^. In response, steps are being taken towards optimizing MRI reliability and increasing sample sizes^8–11^, though this will not be enough. Optimizing biological measurement reliability and increasing sample sizes are necessary but insufficient steps for biomarker discovery; this focus has overlooked the ‘other side of the equation’ - the reliability of clinical and cognitive assessments - which are often suboptimal or unassessed. Through a combination of simulation analysis and empirical studies using neuroimaging data, we demonstrate that the joint reliability of both biological and clinical/cognitive phenotypic measurements must be optimized in order to ensure biomarkers are reproducible and accurate. Even with best-case scenario high reliability neuroimaging measurements and large sample sizes, we show that suboptimal reliability of phenotypic data (i.e., clinical diagnosis, behavioral and cognitive measurements) will continue to impede meaningful biomarker discovery for the field. Improving reliability through development of novel assessments of phenotypic variation is needed, but it is not the sole solution. We emphasize the potential to improve the reliability of established phenotypic methods through aggregation across multiple raters and/or measurements^12–15^, which is becoming increasingly feasible with recent innovations in data acquisition (e.g., web- and smart-phone-based administration, ecological momentary assessment, burst sampling, wearable devices, multimodal recordings)^16–20^. We demonstrate that such aggregation can achieve better biomarker discovery for a fraction of the cost engendered by large-scale samples. Although the current study has been motivated by ongoing developments in neuroimaging, the prioritization of reliable phenotyping will revolutionize neurobiological and clinical endeavors that are focused on brain and behavior.

## Introduction

Biomedical researchers are increasingly recognizing that measurement reliability is a critical determinant of the reproducibility of scientific findings, as it mediates the relationship between sample size, statistical power, and replication between studies^1,2,6,7,21–26^. In response, across a growing number of biological disciplines, researchers are arduously working to optimize the reliability of their assays or tools of choice (e.g., genetics, multimodal MRI, EEG)^3–5,27^. Although of critical importance, these efforts are typically carried out with a singular focus on the biological measurement (e.g., neuroimaging), without ensuring sufficient reliability in the behavioral, cognitive, and clinical (e.g., psychiatric) phenotyping assays commonly employed in studies of brain-behavior relationships. Here, we assert that the lack of focus on optimization of reliability for measures characterizing phenotypic variation is a critical misstep in human neuroscience. This process overlooks the ‘other side of the equation’. It fails to acknowledge that it is the joint reliability (defined as the square root of the intraclass correlation [ICC] of X x Y) of measurements that must be optimized to delineate reproducible brain-behavior relationships.

We draw attention to behavioral, cognitive, and clinical phenotyping as a case in point. While summary constructs for some assessments do show good reliability^28^, a National Institute of Mental Health (NIMH) report outlined that many key cognitive and behavioral phenotypic measures have either not been assessed for their reliability or been found to possess poor to moderate reliability (ICC ≤ 0.4^29,30^). Test-retest reliability in tasks common in cognitive neuroscience are often suboptimal and highly variable, or even incorrectly calculated^31–36^ (i.e., with correlation; e.g., N-back ICC = 0.54, 95% CI = 0.08-0.80; Verbal Memory ICC = 0.46, 95% CI = 0.19-0.64; Attention Network Task ICCs between 0.03-0.66). Furthermore, the most common psychiatric clinical diagnoses have been found to have test-retest reliabilities that are suboptimal for biomarker discovery (Intraclass Reliability [Kappa; K] ≤ 0.7; e.g., schizophrenia K = 0.46; bipolar K = 0.56, borderline personality disorder K = 0.54) with a few being particularly low in field tests of the DSM-5 (e.g., mixed anxiety/depressive disorder K = 0.00; anxiety K = 0.2, depression K = 0.28))^37^. Often regarded as the gold standard for clinical diagnosis, the Structured Clinical Interviews for DSM (SCID) only show good test-retest reliability for depression and specific phobia (2/8; Kappa ≥ 0.7), with moderately better reliability for symptom severity (5/10 diagnoses ICC ≥ 0.7; depression, substance use, post traumatic stress disorder, specific phobia, anxiety)^38^. This has contributed to underwhelming performance and replicability in large-scale neuropsychiatric research^39–42^, as well as to a string of failures in clinical trials and several large pharmaceutical companies moving out of this space^43^. In comparison, MRI data tends to be more reliable than many behavioral measurements or clinical diagnoses, with ICC values between 0.80-0.88 for structural MRI^44^ and up to 0.6-0.8 for more recent ‘optimized’ functional MRI protocols^45^.

Largely, critiques of low reproducibility in neuroimaging studies of individual differences have focused on the inadequacies of MRI data^1,3,46,47^, preprocessing pipelines^8^, and analytic methods^11,48,49^. However, optimizing reliability of one side of the equation (e.g., neuroimaging) without addressing reliability of the other (e.g., phenotyping) leads to underpowered and irreproducible results. This failure to consider the collective impact of phenotypic and biological measure reliabilities is an economically inefficient use of research funding. Additionally, this represents a critical threat to the efficacy of large-sample-focused research, which is likely one of the most promising avenues for the discovery of reproducible brain-behavior associations and biomarkers of mental illness^3–5,50,51^. Remedying this gap is an essential step towards overcoming the reproducibility crisis in psychology and clinical/cognitive neuroscience.

In the current work, we demonstrate that the suboptimal reliability in phenotyping is one of the most significant, and unaddressed, obstacles to biomarker discovery in human neuroscience. Through a combination of experiments on publicly available phenotypic and structural MRI data, simulations, and formal mathematical analyses, we make key takeaway points regarding core issues in biomarker reproducibility. We conclude by offering recommendations for reaching robust conclusions about brain-behavior relationships that are both pertinent to both researchers and funding institutions.

We have also created an interactive statistical exploration tool (Shiny App) available online^52^ to increase the clarity of our findings and to help researchers design well-powered studies. The app will allow readers to examine the complex interactions between study cost, true effect size, estimated effect size, effect size attenuation, statistical power, reliability of brain and behavior measurement, and number of participants and repeated measurements. We offer a guided tour through the Shiny App in the Supplement (S; See S. Figure 1).

## Results and Discussion

### Biomarker discovery depends on joint reliability

In figures A and B, we show that optimizing reliability of one domain (i.e., brain imaging data) without addressing reliability of the other domain (phenotyping) can leave the joint reliability low. This in turn attenuates effect sizes towards zero. We also show that even when neuroimaging ICC is fixed at 1.0, suboptimal phenotypic reliability increases variability in effect size estimates (i.e., correlations) in both normally distributed simulation data (Figure A) and in correlations between structural MRI (sMRI) and IQ data (Figure B). These results drive home the point that even achieving extremely high reliability in one measurement (i.e., neuroimaging) will not enable studies to find effect sizes close to the true effect when the reliability of the other measurement (i.e., phenotyping) is low (e.g., 0.2). We also show that even with moderate joint ICC (0.6), estimated effect sizes can be decreased up to 60% compared to their true effect (S. Figure 2). This happens because suboptimal joint reliability leads to a combination of both downward bias (attenuation) and variability in effect sizes. When joint reliability is low, the estimated effect sizes may often include zero or be negative, which contributes to failures to replicate when these effects are aggregated across studies.

Another important takeaway is that effect size attenuation is maximal for strong effects with low joint reliability (S. Figure 3). All effect sizes are reduced to zero as joint reliability decreases, therefore the size of the attenuation scales with the size of the true effect. Thus, even a very strong effect (e.g., a correlation of 0.9) will be progressively attenuated to zero as joint ICC decreases, meaning that effect size attenuation becomes especially punishing in the search for strong effects. These results emphasize the importance of using multiple raters/measurements or timepoints to increase joint reliability for all effect sizes.

The implications of these findings are of paramount importance. Given the impact of joint reliability on effect size attenuation, we would like to question the acceptance of established brain-behavior effect sizes - as the effects in the literature may be heavily attenuated. Our results show that under an additive error model, the suboptimal joint reliability of current phenotypic and imaging measurements biases brain-behavior effect sizes to zero, making it difficult to assess the statistical validity of brain-behavior relationships. While small and variable estimated effect sizes are highly prevalent in the neuroimaging literature, these effects are strongly attenuated and variable due to imperfect reliability, especially due to suboptimal reliability in phenotyping. In many studies these effects are further compounded by suboptimal reliability in neuroimaging when best practices are not followed^9,11,46,53–57^. We caution readers against interpreting the small effect sizes that are commonly found in large-scale samples^1,58^ as definitive indications of weak or null effects. Furthermore, the replication crisis in neuroimaging is likely due in part to the greater variability in estimated effect sizes that stems from suboptimal joint reliability.

### Effect size attenuation can be corrected using reliability

The impact of effect size attenuation can be corrected if the reliability of data are known (See Supplement). We show in S. Figure 4 that imperfect joint ICC attenuates correlations to zero, but that using joint reliability (i.e., by multiplying estimated correlation by the inverse of joint reliability; see Supplement) corrects these attenuated correlations and can yield effect size estimates that are unbiased by imperfect reliability^9,59^. For example, across both sMRI-IQ correlations and simulated data with moderate effects (r = 0.3) or large effects (r = 0.9), as joint ICC drops, mean estimated effect sizes fall to zero. Notably, correcting for correlation attenuation is most effective for higher ranges of reliability. Lower joint reliability values (< 0.35) show noise in the correction, yielding effect sizes that can be higher or lower than the true effect. However, these corrected values are still much closer to the true effect size than the uncorrected effect size. On the other hand, joint reliabilities above 0.35 tend to yield estimates of effect sizes that consistently reflect the underlying true effect size. This method can be applied to correcting correlation attenuation, though in principle it also applies across other effect size calculations.

### Maximizing joint reliability

In simulated samples of 500 subjects with a true effect of 0.2 (Figure C), we show how the optimization of both sides of measurement reliability is essential in order to reduce both the bias and the variability in estimated effects. The red line shows where effect size attenuation and variability leads to the 95% confidence interval in estimated effects crossing zero, meaning these effects that can no longer be detected consistently across studies. We emphasize joint reliability over individual reliability due to the nature of the joint reliability equation (See Supplemental Methods), which is defined as the square root of the product of ICC_X and ICC_Y. Therefore for any given combination, joint reliability is maximized when both ICCs are equal (i.e., joint reliability when ICC_X = 0.4, ICC_Y = 0.4 > joint reliability when ICC_X = 0.5, ICC_Y = 0.3; See Supplement). For example, to achieve at least a joint ICC of 0.4 when ICC_Y = 0.2, ICC_X must be ≥ 0.8; however, if ICC_Y = 0.4 then ICC_X can be as low as 0.4. With the continuing focus on improving reliability of neuroimaging measurement, it becomes all the more important to assess the extent to which we can remedy the poor reliabilities presented by many phenotypic measures. Recent calls for funding from the NIMH that focus on improving phenotypic reliability highlight the urgency of this need^60^.

### How to improve estimates of reliability

We show in S. Figure 5 that smaller samples yield estimates of reliability (e.g., test-retest, inter-rater) that are highly variable. For example, a study estimating an ICC of 0.4 with only 50 subjects produces estimates of reliability that can vary by over 50%, as is the case even in the DSM-5 field trials^37^. Measures of reliability are subject to a variability proportional to the sample size used in their calculation. For example, an ICC of 0.4 estimated in a larger sample (i.e., n = 500) shows less than 20% variability in the 95% confidence interval. In order to achieve sufficient stability in estimates of reliability, greater sample sizes are required than have been used in most investigations to date.

Aside from increasing sample sizes, inconsistency in estimates of reliability can also be improved by using more than two measurements^61^. As an illustrative example, we calculated the ICC of the Child Behavior Check List (CBCL) and NIH Toolbox items for the longitudinal ABCD study^50^ between the first two years of data acquisition (n = 7,249). We show in S. Figure 6 that the reliability of these variables (ICC = 0.40-0.70) would continue to improve with acquisition of multiple repeated measurements (e.g., four measurements ICC = 0.66-0.91).

Test-retest reliability is also subject to inconsistency as a function of the level of ICC. Lower ICC estimates will have significantly more inconsistency than high ICC estimates. This means that for measures with low reliability, larger samples (n > 500) are required to achieve accurate estimates of ICC. Our Shiny App allows readers to investigate the interactions of these factors in producing accurate estimates of reliability^52^.

These difficulties in accurately estimating reliability prompt the need for large-scale datasets to incorporate repeated measurements (e.g., test-retest, longitudinal data). When large-scale studies collect repeated measurements, joint reliability can be improved, which leads to decreases in effect size attenuation and variability. Furthermore, this practice also enables researchers to accurately measure the reliability of their data, thereby enabling them to correct for the effect size attenuation and recover effect sizes closer to the true effects in question.

### Large samples with low reliability are underpowered

Perhaps most noteworthy is that low reliability in phenotyping makes the collection of large sample sizes ineffective. If the joint reliability of the biological and phenotypic measurements are low, then increasing sample size no longer achieves sufficient statistical power, as demonstrated in Figure D. This shows that if joint reliability is very low (ICC < 0.15), consortium sized samples do not provide sufficient power to detect small-to-moderate effects (r = 0.2), even with five thousand subjects. Conversely, with high joint reliability (ICC > 0.85) even a study with 200 subjects will be better powered to discover reproducible brain-behavior associations than a sample of 5,000 with low reliability.

Moreover, many power calculations (e.g., G*Power) assume the joint reliability of data are perfect, and therefore produce overly optimistic estimates of statistical power^62^. We show here that when statistical power is calculated properly using joint reliability, even moderately sized samples (n ≈ 500) require high joint reliability (>0.6) to achieve sufficient power. When low reliability phenotyping is combined with low reliability brain measurements, such as short resting state scans^46^, or task-based fMRI acquisitions^63,64^, even large-scale samples will not be able to produce robust brain-behavior associations. This finding is consistent with results of recent high-profile work on large-scale samples that have pointed to difficulties in finding consistent brain-behavior associations^9,48,49,51^.

### Repeated measurements achieve higher accuracy

Figure E demonstrates the impact of different sampling strategies on the accuracy of estimated effects as a function of joint ICC. Comparing mean squared error (MSE) between 1,000 subjects with two phenotypic measurements to either 10,000 or 100,000 subjects with only one measurement shows that repeated measurements lead to much lower MSE for only 13% or 1.3% of the cost respectively. Furthermore, increasing sample size beyond a certain point does not significantly impact the accuracy of effect sizes. This effect is driven by the relationship between joint reliability and bias of the effect in question. Bias reflects the average accuracy with which the strength of a brain-behavior relationship is truthfully represented, and bias increases as joint reliability decreases. As a result, increasing joint reliability through aggregating repeated measures yields more accurate estimates of the underlying true brain-behavior relationship. Large-scale (n ≥ 10,000) studies with only a single time-point of phenotyping are a costly and ineffective approach to achieving robust biomarker discovery. We demonstrate that averaging repeated phenotypic measurements can improve reliability^13,14^, though more sophisticated aggregation methods have been shown to perform even better^12,65,66^.

### Large cross-sectional studies are uneconomical

In Figure F, we show that increasing sample size significantly reduces variance in estimated effect sizes, though it does so at a steep price. Increasing sample sizes offers diminishing returns in variance reduction, while increasing study cost linearly. For example, 87% of the reduction in effect size variance achieved from increasing a sample size from 100 to 10,000 is accomplished solely by increasing sample size from 100 to 1,000 participants (Figure F). Assuming a fixed cost of $2000 USD per neuroimaging session and $1000 per clinical and cognitive evaluation, a study with 1,000 subjects with two phenotypic measures can be conducted for only 13% of the cost ($4M USD) of the larger sample ($30M USD; n = 10,000). Critically, this smaller-scale study yields similar variance and much greater accuracy of effect sizes compared to the larger 10,000 subject study.

Importantly, this cost savings becomes even more pronounced as subject sizes increase further to 100,000 subjects and beyond. Recent estimates have shown that large samples (n > 2,000) are required to discover robust brain-wide associations with behavior^1^. Our results demonstrate similar effects, showing how large sample sizes are helpful, but up to a point. Prioritizing ever larger samples without improving measurement reliability becomes increasingly less helpful for establishing brain-behavior relationships that are both reproducible and accurate. We demonstrate that more moderately sized samples (n = 1,000) with aggregated repeated measurements will both save time and cost while leading to significant increases in robust biomarker discovery through improved phenotypic measurement reliability.

### When to prioritize reliability versus sample size

S.Figure 7 shows that for every point in the possible three-dimensional space of sample size, joint ICC, and effect size, that there is an optimal study design choice that will lead to the largest reductions in MSE. For smaller studies, it tends to be most effective to increase sample size, but as samples become larger (n > 500), improving joint reliability starts to lead to relatively greater decreases in MSE compared to increasing sample sizes. This demonstrates that for larger samples, improving measurement reliability through repeated measurements become especially helpful in producing robust estimates of brain-behavior associations. Importantly, this holds regardless of the effect size or joint reliability in question. For strong effects (r ≥ 0.7), increasing joint reliability always leads to greater reductions in MSE than increasing sample sizes. We provide an interactive version of this plot with a path-finding function in the Shiny app to help researchers explore the possibility-space of these choices in the design of their study and how they interact with MSE, variance, and statistical power.

S. Figure 8 demonstrates the variability in effect sizes as a function of sample size and joint ICC. Increasing sample sizes decreases the range of the upper and lower bounds of estimated effect sizes, with most of the decrease in variability coming from increasing samples from 0 to 500 subjects. Notably, prioritizing even larger sample sizes when joint reliability is low (< 0.2) can still produce effect sizes with lower bounds that cross zero, indicating these effects are not likely to reproduce across studies. These results demonstrate the consequence of optimizing joint reliability of measurement, as for most effect sizes and moderately large samples (n > 500), the best way to improve the accuracy and replicability of a study is by increasing joint reliability.

## Recommendations for Researchers and Funders

Discouragingly small estimates of brain-behavior effect sizes and the large samples needed to detect them have grown increasingly concerning^67^. A key question is: can the points made here help change the picture? We believe the current work demonstrates that the optimization of reliable phenotypic measures, in a fashion similar to what has been labored over in the neuroimaging community^3–5,11^, opens up the potential to appreciate larger and more reproducible effects for brain-behavior relationships than reported to date. We provide suggestions for researchers and funding bodies when considering new research projects to maximize the likelihood of biomarker discovery.

### Use reliability as a guide

Researchers should use inter-rater and test-retest estimates of reliability to evaluate if their variables of interest are reliable enough to find reproducible brain-behavior relationships^9,46,68^. The results demonstrated here on ICC can be generalized to other measurements of reliability (e.g., Kappa) given that the underlying generative function is similar. Prior work has demonstrated comparable closed form solutions for ICC and weighted Kappa^69^. If a study aims to examine phenotypic variables known for low reliability (e.g., DSM depression diagnosis), it is important to not only acquire repeated phenotypic measurements or multiple raters, but moreover to focus on brain measurements with the highest reliability (i.e., structural measures or long acquisition functional connectivity [>20 minutes]). Conversely, for a study looking for brain-behavior associations with neuroimaging measurements that have low reliability, use only the most reliable phenotypic measures possible (e.g., age, sex, IQ, etc), and not cognitive or clinical measures with suboptimal reliability.

### Do not pursue statistical validity over reliability

Reliability places an upper bound on statistical validity (i.e., effect sizes). As such, any effort to maximize statistical validity would benefit from consideration of reliability. Statistical validity of brain-behavior associations cannot be assessed or prioritized accurately or reliably without achieving sufficient measurement reliability first. In low reliability scenarios, any estimates of statistical validity (e.g. brain-behavior correlation) will be highly variable and attenuated close to zero (Fig A & B). Estimates of statistical validity are subject to noise, and studies prioritizing strength of statistical validity over reliability must contend with both inaccurate and variable estimates of effect sizes as a result of suboptimal reliability. This means that in one study effect sizes may appear large, but small in another study, impeding the ability to find consistent strong effects across studies and inducing failures to replicate. Optimizing for reliability alone is also insufficient however, as sources of noise can be highly reliable and will only serve to reduce interpretational validity, even if they improve reliability and statistical validity^3–5^. For example, head motion in fMRI corrupts the interpretational validity of fMRI connectivity while being both highly reliable^70^ and sensitive to clinical presentation^71^ and associated brain-behavior relationships^72^.

Regardless of the considerations of optimizing study design for either reliability or statistical validity, any study should include assessments of reliability to improve their ability to detect effects. Incorporating measurement reliability into efforts to optimize effect sizes enables correction for effect size attenuation, which is a significant issue in clinical and cognitive neuroscience, and is most severe for large effect sizes.

### Prioritize repeated measurements more than large samples

Collecting multiple raters/measurements/time points per participant is often a more economical method for maximizing scientific reproducibility than increasing sample size (Figure E, F)^12–15^. When possible, we recommend using multiple clinicians and/or repeated assays to evaluate clinical presentation and cognition. Given the challenges of collecting repeated measurements, recent advances in data collection strategies may enable such repeated assessments outside the laboratory setting (e.g., ecological momentary assessment and cognitive burst sampling via smartphone- or web-based assessments). These approaches can be leveraged to mitigate experimenter and participant burden, as well as to increase the accessibility of participation and reduce participant dropout^16–20^.

### Use the provided Shiny app

Plan out studies regarding expected ICC, true effect size, and sample size using our Shiny App. A study is likely to replicate when the lower bound of the 95% confidence interval (S. Figure 1; Panel A) in estimated correlations falls in the direction of the expected true effect.

## Conclusions

In the current work we offer perspectives on how suboptimal behavioral, clinical, and cognitive phenotypic reliability hinders biomarker discovery through its interaction with sample size and estimated effect sizes in brain-behavior relationships. We have shown that optimizing joint reliability must be prioritized to improve the accuracy and reproducibility of our estimated brain-behavior correlations. Using more reliable measures allows for robust estimates of true effects even at smaller sample sizes, and repeated measurements can be leveraged to provide more accurate estimates of brain-behavior relationships with one tenth the sample size and for a fraction of the cost. We expect the impact of aggregating repeated measurements to improve reliability will hold under most conditions, though the exact relationship or performance observed may differ under more complicated error structures. We hope that the perspective shared here will inform the design of new studies and the analysis of already collected data.

**Figure 1.**
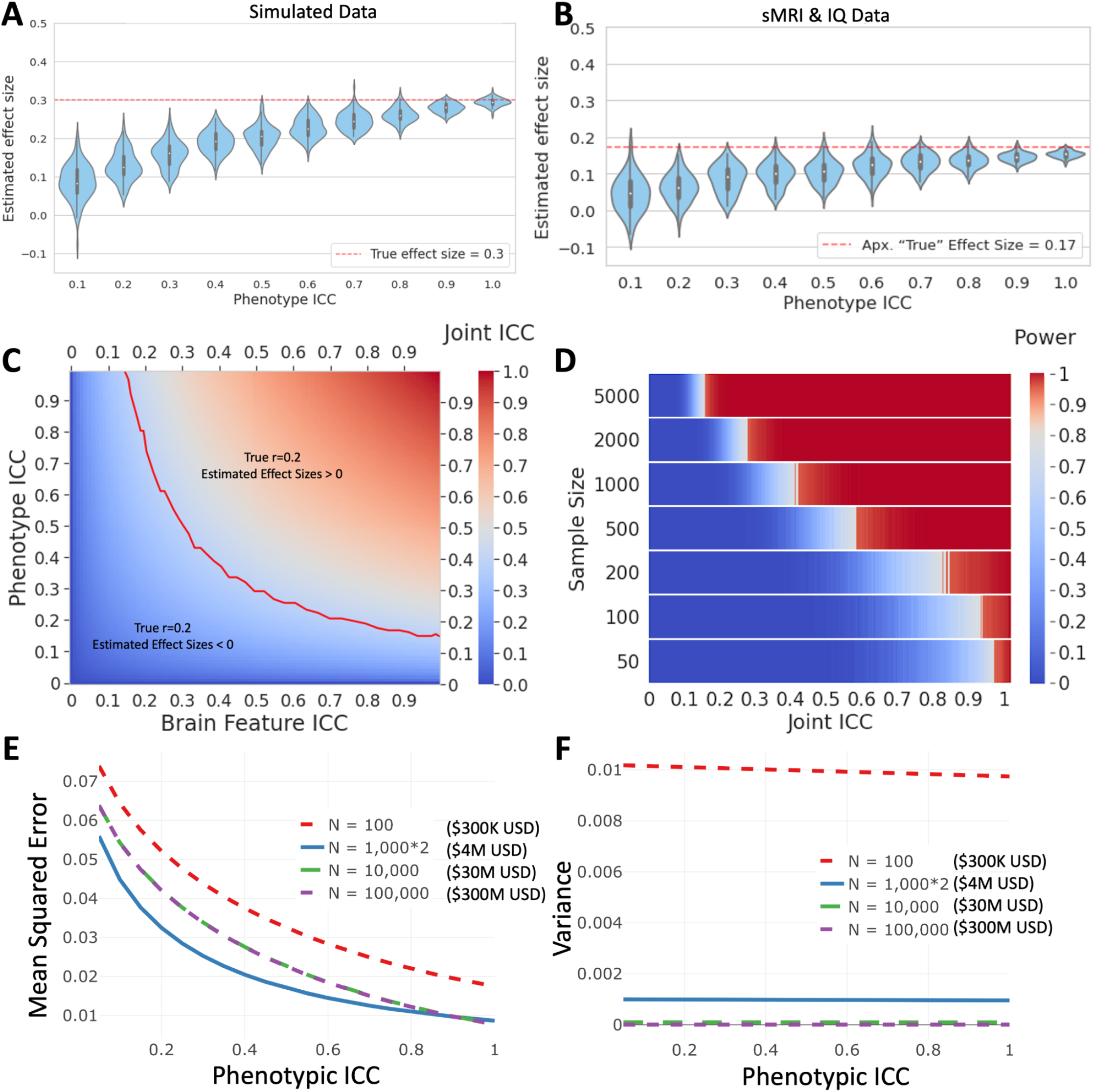
A) We show the distribution of estimated effect sizes in simulation data with 500 subjects and a true brain-behavior correlation of 0.3. Even when ICC_X (brain measurement) is perfect (ICC = 1.0), lack of phenotypic reliability significantly attenuates the estimated correlation to zero. This demonstrates that suboptimal joint reliability is an important cause of lack of reproducibility in neuroimaging studies and reason why large-scale samples often demonstrate very small effect sizes. B) We perform the same calculation as in A, but with synthetic data created from real effect sizes and distributions taken from two structural scans in 568 individuals associating cortical thickness measures with intelligence. These results confirm that the results of the simulations hold in real data. C) This heatmap shows the relationship between joint ICC and the attenuation and variability in effect sizes (n = 500). As joint ICC increases, effect size attenuation and variability decrease, meaning estimated effect sizes will become both more accurate and reproducible from study to study. The red line shows the minimum ICC required for phenotyping and brain measurements to be able to consistently estimate a positive relationship across studies when the true underlying correlation = 0.2. D) This heatmap shows the relationship between statistical power and joint reliability (ICC) of brain measurement and phenotypic data as a function of sample size with a true effect of 0.2. When sample sizes are exceedingly low (n < 200), only the most reliable data will be well powered to discover brain-behavior correlations. As joint reliability drops however, even large-scale samples are no longer well-powered. When joint reliability is low (ICC < 0.2), even samples with thousands of participants are not sufficiently powered to recover true brain-behavior correlations. E) We investigate the power of repeated phenotypic measurements to increase reliability and improve the accuracy of brain-behavior correlations as a function of joint ICC (fixed neuroimaging ICC = 0.5). The red, blue, green, and purple lines correspond to the average accuracy (MSE) of samples with 100 subjects (1x neuroimaging/phenotyping), 1000 subjects (1x neuroimaging, 2x phenotyping), 10,000 subjects (1x neuroimaging/phenotyping), and 100,000 subjects (1x neuroimaging/phenotyping). Assuming a fixed cost of $2000 per neuroimaging session and $1000 for clinical and cognitive phenotypic evaluation the study design with 1,000 subjects measured twice is able to achieve better accuracy for $4M USD, a fraction of the cost of larger samples ($30M USD, n = 10,000; $300M USD, n = 100,000). Notably, increasing sample size from 10,000 to 100,000 does not yield a notable increase in mean accuracy. F) The primary value of acquiring large sample sizes lies in the decrease in variance of estimated effect sizes. We compare the variance in estimated effect sizes as a function of joint ICC and different sampling strategies. Increasing samples from 100 subjects measured once to 10,000 subjects measured once yields a large decrease in variance in estimated effects. However, most of the gain in variance reduction happens in the first 1,000 subjects. 87.5% of the variance reduction in moving from 100 to 10,000 subjects is achieved by 1,000 subjects with phenotyping measured twice, for less than a seventh of the total cost. Increasing sample size from 10,000 to 100,000 yields only a small reduction in variance in effect sizes for 10x the cost. This points to the fact that increasing sample sizes impacts the variance in estimated effect sizes logarithmically, while study costs continue to increase linearly. A much more cost effective approach would be to measure subjects twice to increase the reliability of the measurements, thereby achieving most of the variance reduction while also significantly increasing accuracy of estimated brain-behavior associations.

## Data Availability

All data used in the current work are available through the Healthy Brain Network, an open source resource for transdiagnostic mental health research: http://fcon_1000.projects.nitrc.org/indi/cmi_healthy_brain_network/

## Code Availability

We provide an open source Shiny App for researchers to evaluate our results: https://andrew-a-chen.shinyapps.io/reliability-app/. All additional codes in the current work will be made publicly available upon publication of the manuscript.

## Methods

### Overview

We evaluate the impact of low phenotypic reliability primarily through simulation data and synthetic data derived from publicly available structural MRI and IQ data to examine: 1) the impact on the power, accuracy, and variability of brain-behavior associations from increased reliability versus increased sample size; the bias in correlation estimation with unreliable data at a given sample size, and in our Shiny App^52^, we show how estimates of reliability can be used to correct for this attenuation; 3) the impact of collecting multiple measurements per individual to improve reliability; 4) the relative cost and likelihood of accurate effect size estimation (MSE and variance) of studies with different sample sizes and numbers of measurements per person.

### MRI and Phenotypic Data Acquisition

The sample included 568 children and adolescents from the Healthy Brain Network cohort^73^ with 364 males and 204 females. All participants were between ages 6-17 (mean 10.45, SD 2.69). Participants’ IQ (mean 102.34, SD 17.36) was measured using the Wechsler Intelligence Scale for Children (WISC-V)^74^. Participants were recruited on a community self-referred basis through the distribution of advertisements and announcements to community members, educators, local care providers, and parents. Main exclusion criteria included the presence of acute safety concerns (e.g., danger to self or others), cognitive or behavioral impairments that could interfere with participation (e.g., being non-verbal, IQ less than 66), or medical concerns that are expected to confound brain-related findings^73^. Anatomical MRI scans were acquired for all participants using both standard HCP T1w and VNAV T1w MPRAGE. Acquisitions for all participants were obtained at a single site from the Cornell Brain Imaging Center (CBIC) using a 3 T Siemens Prisma scanner. The full set of T1 image acquisition parameters can be found in the HBN data release documentation^73,75^

### MRI Preprocessing

The skull-stripped anatomical images and raw functional images were preprocessed through the Configurable Pipeline for Connectomes (C-PAC^76,77^. Anatomical images were nonlinearly registered to the MNI152 template^78^ (2 mm isotropic) using Freesurfer^79,80^ and segmented into gray matter (probability threshold = 0.95), white matter (probability threshold = 0.95) and cerebrospinal fluid (CSF; probability threshold = 0.95). We used Mindboggle^81^ to extract a total of 70 cortical thickness values from the left and right hemisphere (35 features per hemisphere). Mindboggle achieves high accuracy gray matter extraction by merging structural mesh models from both Freesurfer and ANTs^82^. We extracted cortical thickness values for both HCP-T1 and VNAV T1 MPRAGE images. Cortical thickness values from HCP-T1 and VNAV T1 were then harmonized to correct for acquisition differences using a version of ComBat specialized for longitudinal data^83^.

### MRI Analysis

We avoided univariate analysis between structural features and IQ to prevent overfitting, opting for a multivariate dimensional approach. We performed Partial Least Squares (PLS) on the 70 cortical thickness values, to identify the primary dimensions of structural variance associated with IQ. PLS, similar to principal component analysis (PCA), creates components that are a linear combination of each of the cortical thickness values that maximize their covariance with IQ. PLS was performed separately on harmonized HCP-T1 and VNAV T1 features. All PLS components were then matched based on correlation across participants, and the component that showed high alignment between HCP and VNAV T1 scans and strongest average association with IQ (r = 0.175) was selected for further MRI-IQ analysis and synthetic data analysis.

### Synthetic and Simulation Data Analysis

Using both real data (MRI, IQ) and simulated data, we generated correlated datasets from bivariate normal distributions with varying amounts of added measurement errors to obtain variables with desired ICC levels (details in Supplementary section, “Simulation Setup”). In brief, we generated simulated brain imaging (X) and phenotypic variables (Y) based on an additive error model. Across all simulated sample sizes (50-10,000) and true correlation values (0.1-0.9), noise free X and Y are generated with a fixed true correlation. We manipulate the ICC by adding variance to the noise term. After calculating variances, we simulate noise for X and Y 1,000 times and this gives an X and Y pairing for each ICC value, sample size, and true correlation value. We calculate the upper and lower confidence intervals and mean estimated correlation. This process is then repeated 100 times and the average of upper and lower confidence intervals for estimated correlation are calculated. Using these confidence intervals we can then calculate power at each given ICC value for the brain (ICC_X) and behavior (ICC_Y) measurements. Under each scenario, we calculate correlations with and without attenuation correction using known ICC (See Shiny App). We performed the same process for synthetic data analysis, using the real sMRI and IQ values and covariances to form our synthetic data. PLS components were extracted from both T1 images and matched based on correlation across participants (r>0.9). The average between component scores correlation with IQ was regarded as the approximated “True” effect size (r = 0.175). The VNAV scan was considered an approximated “True” measurement in order to synthetically create a perfectly reliable estimate of the PLS component. We corrupted this component with Gaussian noise in a stepwise fashion to create brain measurements across several levels of ICC.

## Supplementary Methods

### 1. Interactive visualizations of simulation and theoretical results

We provide a set of R Shiny-based interactive visualizations for researchers to explore our main results.

**Supplementary Figure 1** shows screenshots of each tab in the shiny app, which we describe briefly below:

#### Simulation results across ICC values (A)

We examine how varying degrees of reliability impacts correlation estimation, shown through several evaluation metrics using both simulated and real data. 3D surface plots are used to display various simulation results, including the mean correlation estimates and variability of those estimates. Data for the correlation between structural MRI and IQ is from the Healthy Brain Network cohort, with details available in **Methods**. For real data, normally distributed measurement error is used to simulate varying ICC of fMRI and IQ data.

#### Theoretical results for averaging repeated measures (B)

We assess theoretically, what are the relative benefits of increasing sample size using a single measurement vs. using repeated measures in a smaller sample. We use line plots to display the theoretical benefits of averaging repeated measures in a subsample over using a larger sample with single measurements. Points at which ICC estimation in the subsample with repeated measurements has a lower MSE or lower variance than the full sample are displayed across graphs. Users can change several parameters including sample size, number of repeated measures, ICC, and true correlation.

#### Increasing sample size versus collecting another repeated measure (C)

This plot helps readers evaluate for a given correlation, reliability, and sample size: is it more advantageous to increase sample size or collect a repeated measure for each subject? Using simulation results, cone plots provide guidance on whether to increase sample size or collect more repeated measures for various outcome measures of interest. The direction of the cone reflects whether the outcome measure increases more if a researcher collects more samples or improves reliability by 0.1 through collecting repeated measures. Clicking on a cone will draw a line toward the directions of maximal benefit. The size of each cone represents the relative size of the benefit in the direction of the cone.

#### Deflation of correlation (D)

This plot examines how reliable measurements need to be in order for estimated correlations to stay above a designated threshold. The left of each line on the plot is the region of values at which the estimated correlation in simulation drops below a specified threshold. True correlation values are displayed as different colors. Options are provided to vary the sample size and also to view the results across all sample sizes in the simulation.

#### Accuracy of ICC estimation (E)

We examine the uncertainty of ICC estimates over varying sample sizes. Plots are used to show the accuracy of intraclass correlation estimation across sample size and the proportion of subjects with repeated measures. Results are shown from the main simulations, where the chosen proportion of subjects have a single repeated measure.

In real data examples, ICC themselves are often estimated with a small subset of the samples with repetitions. Here we demonstrate the uncertainty associated with estimating an empirical ICCs based on the subset of the full samples. Line plots are used to show the accuracy of intraclass correlation estimation across sample size and the proportion of subjects with repeated measures. Results are shown from the main simulations, where the chosen proportion of subjects have a single repeated measure.

**S. Figure 1.**
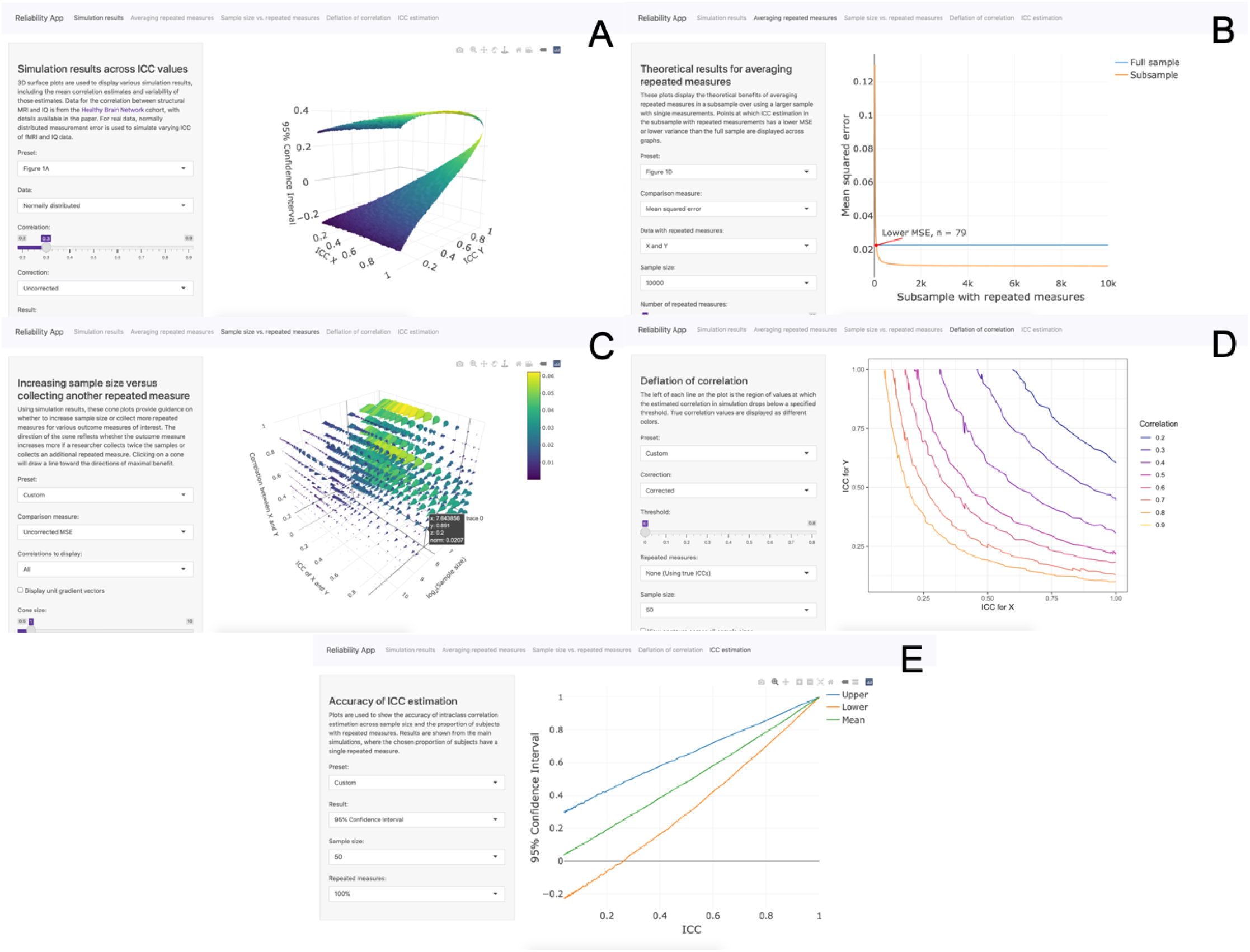
Pictured above are each of the tabs of the Shiny App as explained above.

### 2. Simulation setups

We design simulations to assess the impact of reliability and effectiveness of the attenuation correction across a broad range of parameters. Let measurements without error *A*_*i*_ and *B*_*i*_, *i* = 1, 2, …, *n* be drawn from correlated normal distributions with variances σ^2^ = 1 and correlation ρ. Then we generate measurements observed with error via

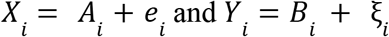

where *e*_*i*_ and ξ_*i*_ are Gaussian noise with mean 0 and variance σ_*e*_^2^ and σ ξ^2^ respectively. By varying σ_*e*_^2^ and σ_ξ_^2^, we simulate data with varying levels of reliability, measured through the ICC values

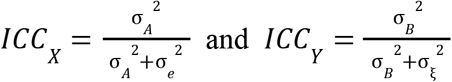

#### Using true ICC

We first compare the estimated correlation with and without the correction, assuming we know the underlying ICC values *ICC*_*X*_ and *ICC*_*Y*_. We first sample simulated observations without error *A*_*i*_ and *B*_*i*_ then for every set of ICC values, we draw 1,000 sets of measurements *X*_*i*_ and *Y*_*i*_, *i* = 1, 2, …, *n* from the proposed model and then calculate the uncorrected correlation *r*_*X,Y*_ and corrected correlation 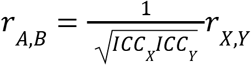 for each set. Across these 1,000 draws, we compute key summary statistics including the mean, variance, and 95% confidence interval (2.5th and 97.5th percentiles). We then repeat these steps for 100 random draws of *A*_*i*_ and *B*_*i*_ and report the average of those summary statistics.

#### Estimating ICCs from repeated measures

To account for uncertainty in ICC estimation, we extend the simulation to include repeated draws for a proportion *p* of the observations under the same measurement error model

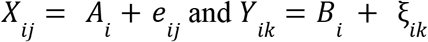

where *i* = 1, 2, …, *np, j* = 1, 2 and *e*_*ij*_ and ξ_*ik*_ are Gaussian noise with mean 0 and variance σ_*e*_^2^ and σ_ξ_^2^. For each of the 1,000 sets of measurements with error, we obtain ICC estimates 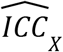 and 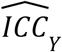 from these observations with repeated measures and report the mean, variance, and 95% confidence interval of these estimates. We now use these estimated ICC values to calculate the corrected correlation as 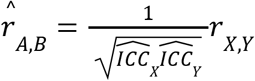 and again report summary statistics, averaged across 100 random draws of *A*_*i*_ and *B*_*i*_.

#### Synthetic data setup using Healthy Brain Network sMRI and IQ data

We modify our simulation setup to incorporate data from the Healthy Brain Network cohort to show how measurement error can impact the estimated correlation between structural MRI and IQ. Instead of simulating *A*_*i*_ and *B*_*i*_, we repeat our simulations using the top PLS component from the cortical thickness features (see **Methods**) as *A*_*i*_ and IQ as *B*_*i*_ where *i* = 1, 2, …, *n*. Individual simulations are conducted by sampling *n* subjects from the full 568 HBN observations with replacement. The subsequent simulation steps are the same as previously outlined.

#### Software

All analyses are performed using R version 4.1.1. ICCs are calculated using the *psych* package (Version 2.1.9) using ICC1 from the *ICC* function.

### 3. Measurement error and attenuation bias in estimating correlations

Measurement error and its impact has been commonly studied in the statistical literature. For the problem of our concern, we suppose X’s are brain signatures (e.g., functional connectivity metrics) and Y’s are clinical or cognitive phenotypic measurements. The observed data are measured repeatedly with noise. For each individual, we assume a simple additive measurement error model as

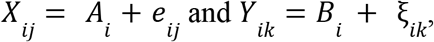

where *e*_*ij*_ and ξ_*ik*_ are random noise with mean 0 and variance σ_*e*_^2^ and σ_ξ_^2^, respectively. The population ICCs for each of the variables can be defined as,

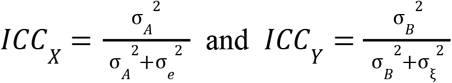

The population Pearson’s correlation with a single measurement of X and Y could be then written as

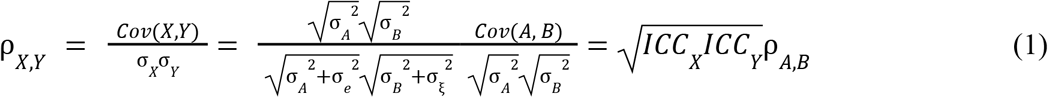

Since 0 ≤ *ICC* ≤ 1, the observed Pearson correlation tends to underestimate the correlation of the underlying features by an attenuation factor equal to the joint reliability of 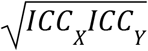.

Similar relationship holds for the empirical sample version of the correlations, as denoted by *r*_*X,Y*_ and *r*_*A,B*_.

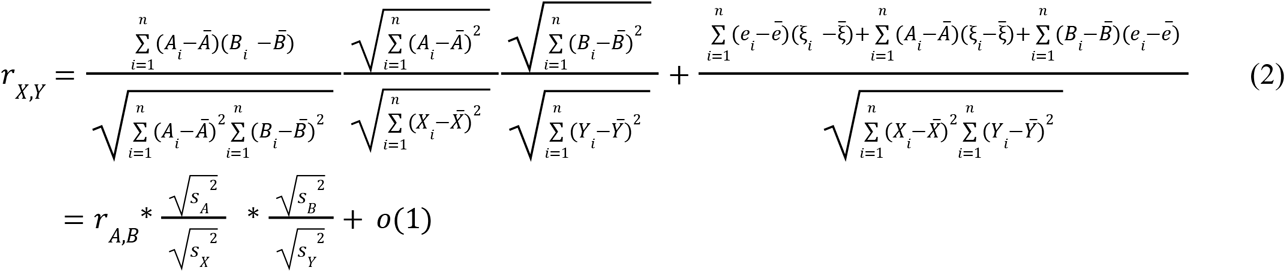

#### Correction of Pearson’s correlation by attenuation factor

Based on (1) and (2), it becomes straightforward that if there exist external estimates of the corresponding ICCs, we could apply a correction to adjust for the bias in estimating ρ_*A,B*_ via:

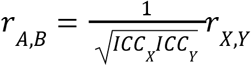

We could approximate the variance as

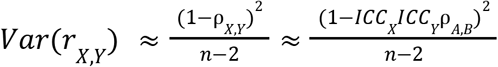

This means that, the lower the ICCs are for each of the measures, the higher the variability is in the obtained correlations between X and Y. If corrected values exceed an absolute value of 1, these corrected correlations can be constrained to −1 to 1. Corrected correlation exceeding an absolute value of 1 is likely due to variance in the estimated correlation. The variance of the ICC-corrected correlations between A and B can then be approximated as:

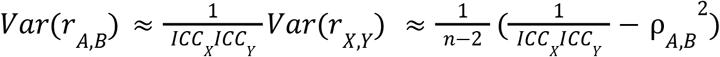

Still the lower the ICCs are, the larger the variances are. So far we haven’t accounted for the uncertainty in estimating ICCs as we assume that those could be consistently estimated using external datasets with large enough sample size.

#### Trade-offs between repeated measures and large samples with a single observation

We now investigate the gains and costs in terms of estimation bias and variance when we have repeated measures over relatively smaller samples versus when we have a large number of subjects with a single observation. Denote this correlation 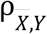 where 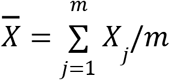 is the average of *m* repeated measures. Under the assumption that measurement errors are independent of each other and *A*_*i*_, it has variance

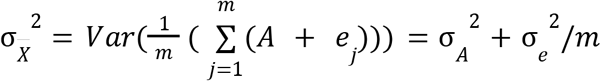

Then the correlation with clinical or cognitive phenotypic measurements becomes:

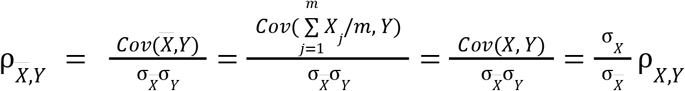

Denote 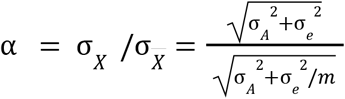. Note that as *m* increases, α approaches 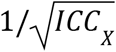, so 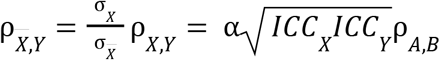 approaches 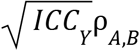 and the deflation of ρ_*A,B*_ due to measurement error in *X* becomes increasingly negligible. The variance of the Pearson correlation estimate using the averaged values 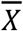 is now

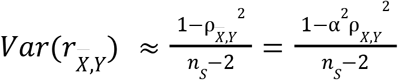

Now, we aim to compare this to the variance of the Pearson correlation estimate across all subjects using only their single measurement. The ratio between the variance of these two alternative estimates is

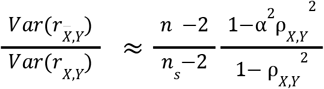

Which depends on *n, n*_*S*_, α, *and* ρ_*X,Y*_.

We can also compare these estimates via mean squared error (MSE) where the bias is calculated with respect to the correlation between *A* and *B* (without measurement error)

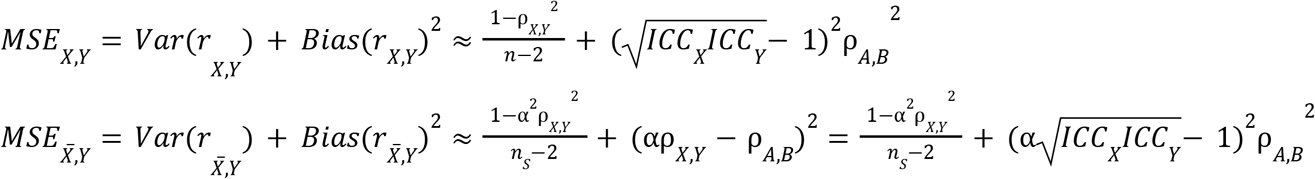

which can then be compared via their difference.

Similarly, suppose we also have *p* repeated measures for *Y* and aim to evaluate the performance of the averaged measurements 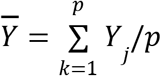. We then find that

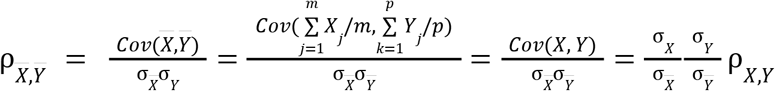

since 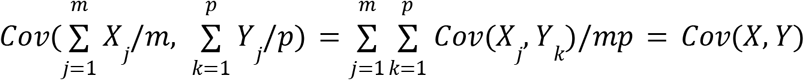.

#### Improvement in population ICC by averaging repeated measures

Under the additive measurement error model, we can derive the population ICC of *m* averaged measurements in terms of the original population ICC. Let *X*_*ij*_ denote repeated measures where *i* = 1, 2, …, *n* indexes subjects and *j* = 1, 2, …, *m* indexes the number of repeated measures, which is assumed to be constant across subjects. We assume a simple additive measurement error model

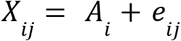

Where *A*_*i*_ are random variables with variance σ_*A*_^2^ representing the value measured without error and *e*_*i j*_ is random noise with mean 0 and variance σ_*e*_^2^. The population ICC of *X*_*ij*_ is defined as 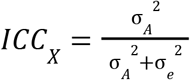.

Denote the random variables obtained by averaging the *m* repeated measures as 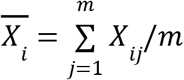. Based on the derivation from the previous section, the corresponding population ICC of the random variable 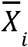 is given by

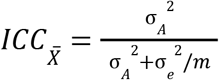

Using the fact that the error variance σ_*e*_^2^ can be written in terms of σ_*A*_^2^ and *ICC*_*X*_ as σ_*e*_^2^ = σ_*A*_^2^/*ICC*_*X*_ − σ_*A*_^2^, we can rewrite the population ICC for the averaged measurements as

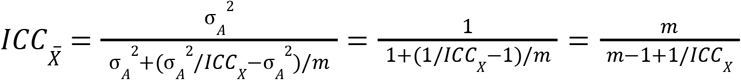

## Supplementary Results

**S. Figure 2.**
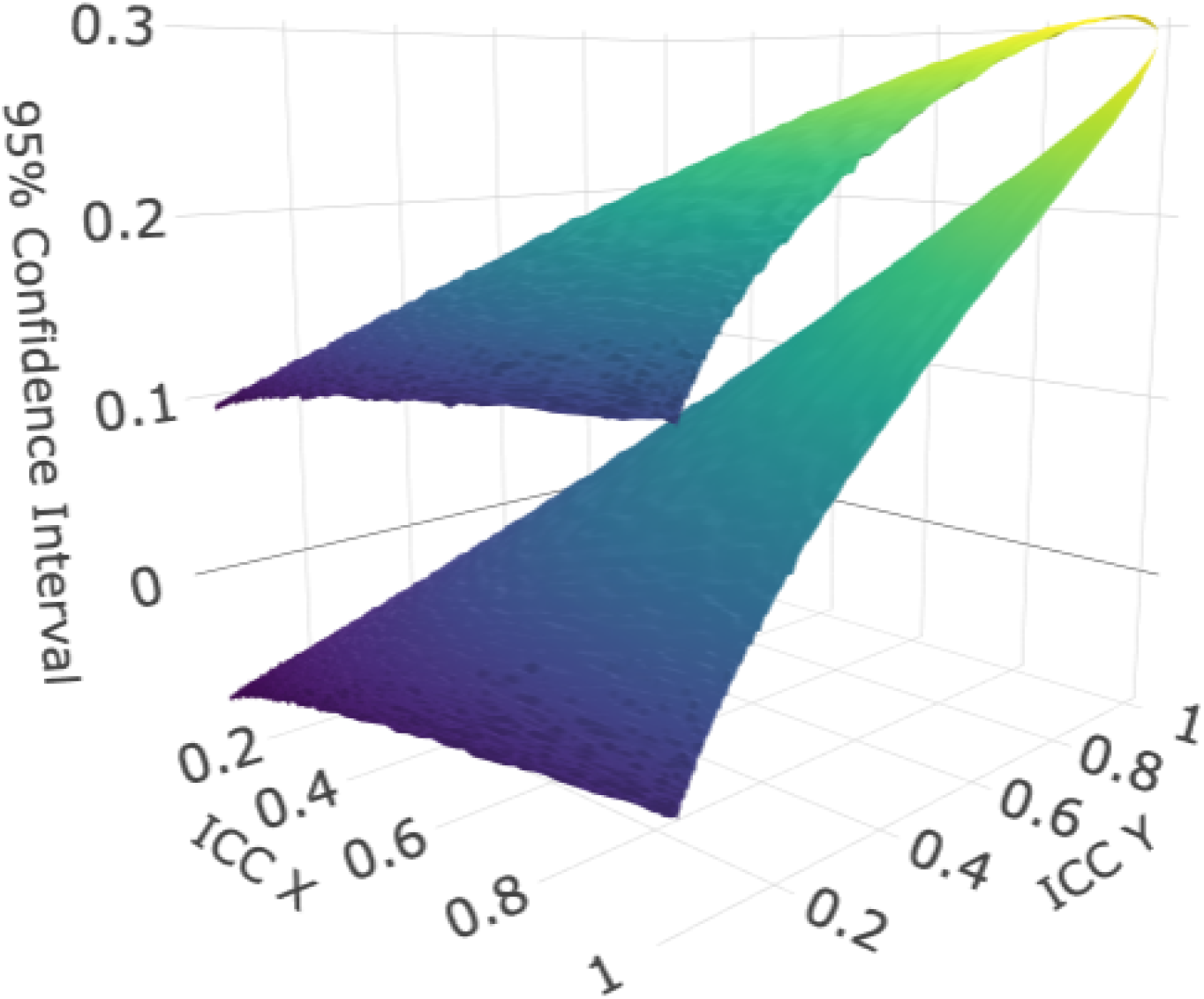
The 95% confidence interval of estimated effect sizes as a function of the ICC of X and Y. Upper and lower bound correspond to the expected variability in effect sizes for a given joint ICC level. Sample size of 500, and a true effect size of r = 0.3 are used for this simulation. High ICC for both X and Y (>0.8) prevents effect size attenuation. For an average joint ICC that may be observed in an imaging study (ICC X = 0.6, ICC Y = 0.6), the lower bound correlation = 0.1, showing that even moderate reliability (ICC = 0.6) can suffer from effect size reduction of up to ∼ 60%.

**S. Figure 3.**
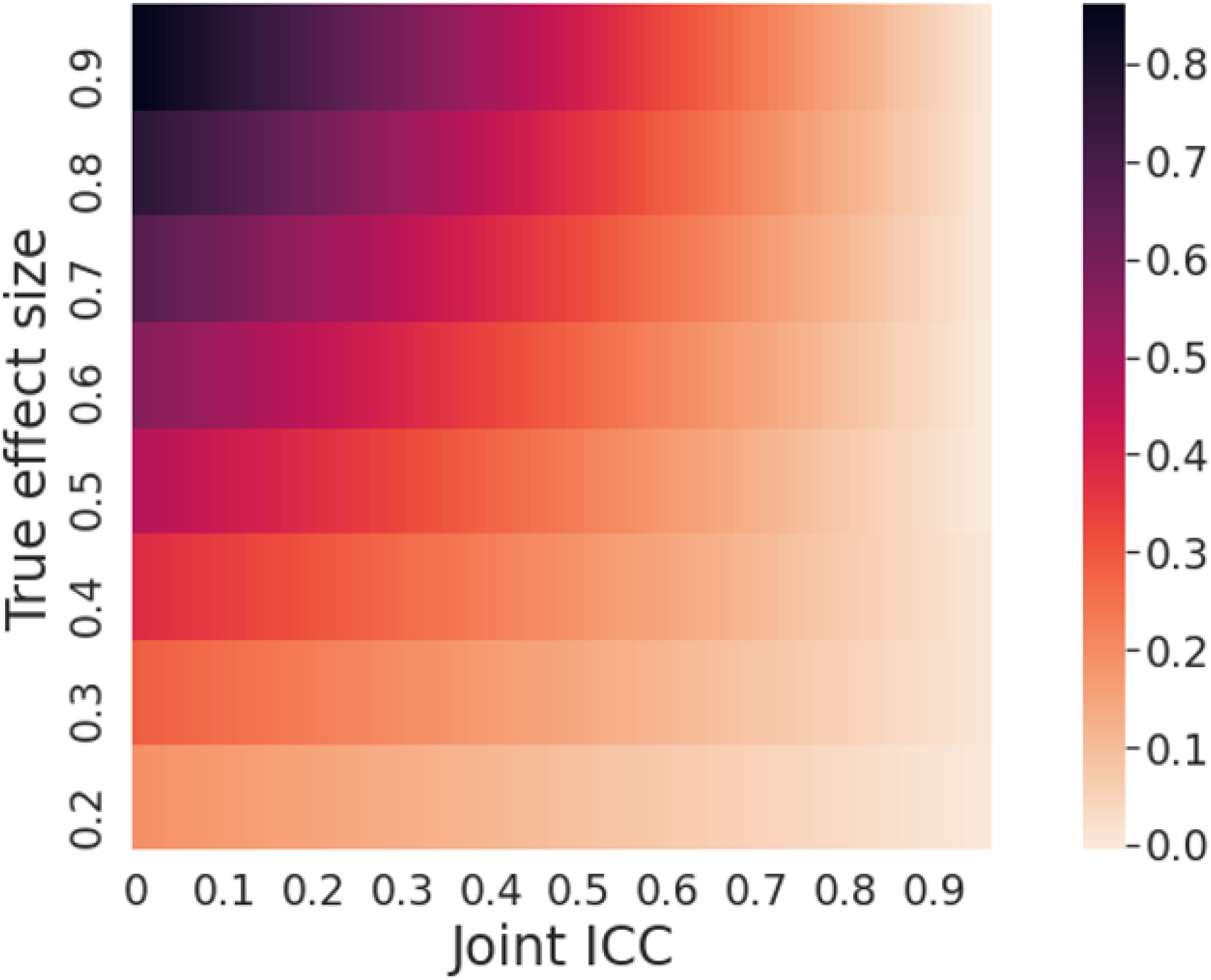
We summarize the severity of effect size attenuation as a function of Joint ICC and the true underlying effect size. As Joint ICC decreases, attenuation increases, resulting in larger differences between the average estimated effect size and the true effect size. Absolute level of attenuation also increases as a function of the true underlying effect size, meaning that stronger effects will be attenuated more than weak effects. The largest attenuation can be seen for strong effects where X and Y have poor joint reliability, as even the strongest effect sizes will be attenuated to zero.

**S. Figure 4.**
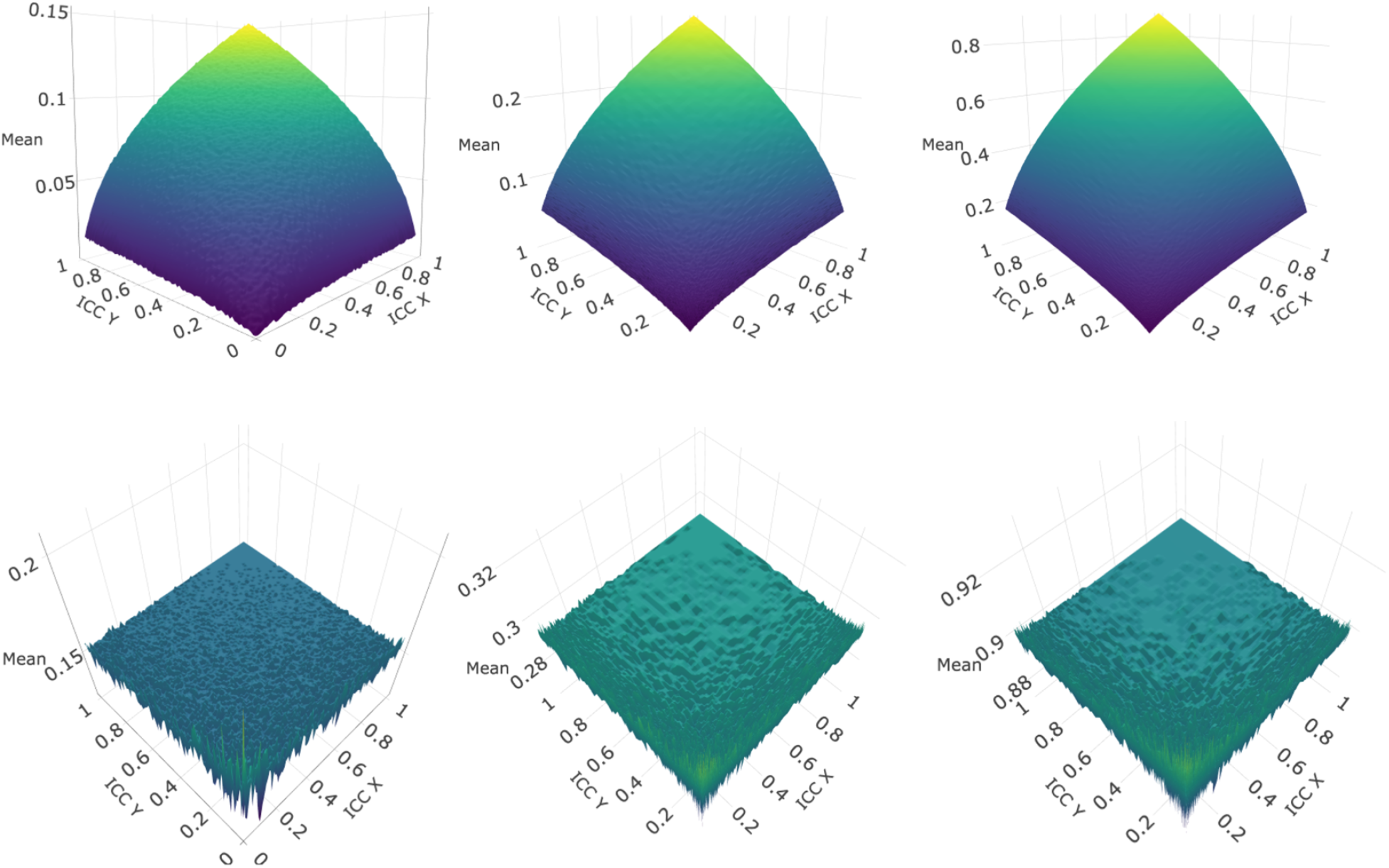
The top row shows the attenuation of average estimated correlations at different levels of ICC_X and ICC_Y. The bottom row shows the average estimated correlations for different levels of ICC_X & ICC_Y after ICC-correction has been applied to remove attenuation effects. First, second, and third columns correspond to MRI-IQ data (n = 500), simulation data with r = 0.3 (n = 500), and simulation data with r = 0.9 (n = 500), respectively. Higher ICC results in more accurate corrections of attenuation, though even corrections with low reliability yield estimates that are more accurate than the uncorrected effect sizes. Taken together, this shows that any effect size can be attenuated to zero, but that this attenuation can be corrected in the reliability of X & Y have been calculated.

**S. Figure 5.**
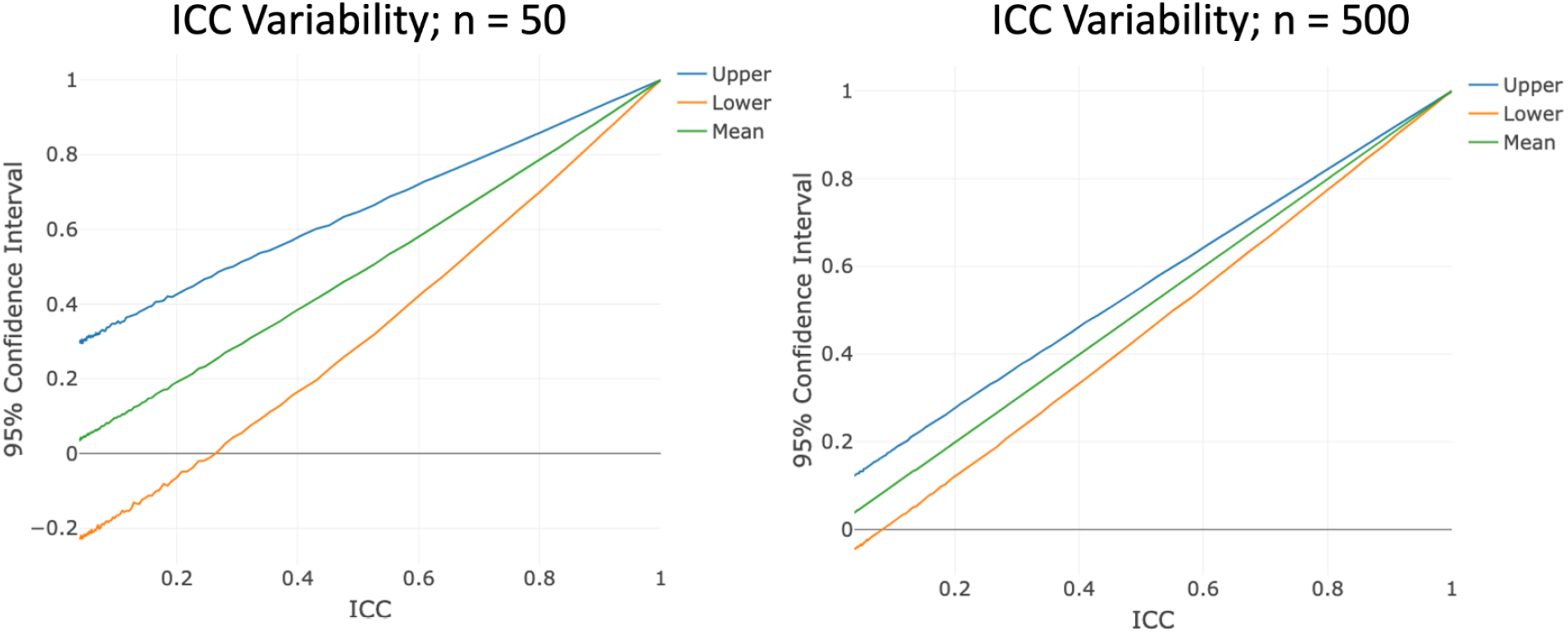
Variability in ICC estimation depends on both the ICC and the number of subjects. As ICC increases, variability in the ICC estimation decreases. In other words, higher ICC values will on average be more accurate representations of the true underlying ICC. Small samples create variable estimates of ICC, in A we show the True ICC and variability in observed ICC as a function of ICC level for a sample of 50 participants. B shows the variability for 500 subjects, and shows much lower levels of variance compared to A.

**S. Figure 6.**
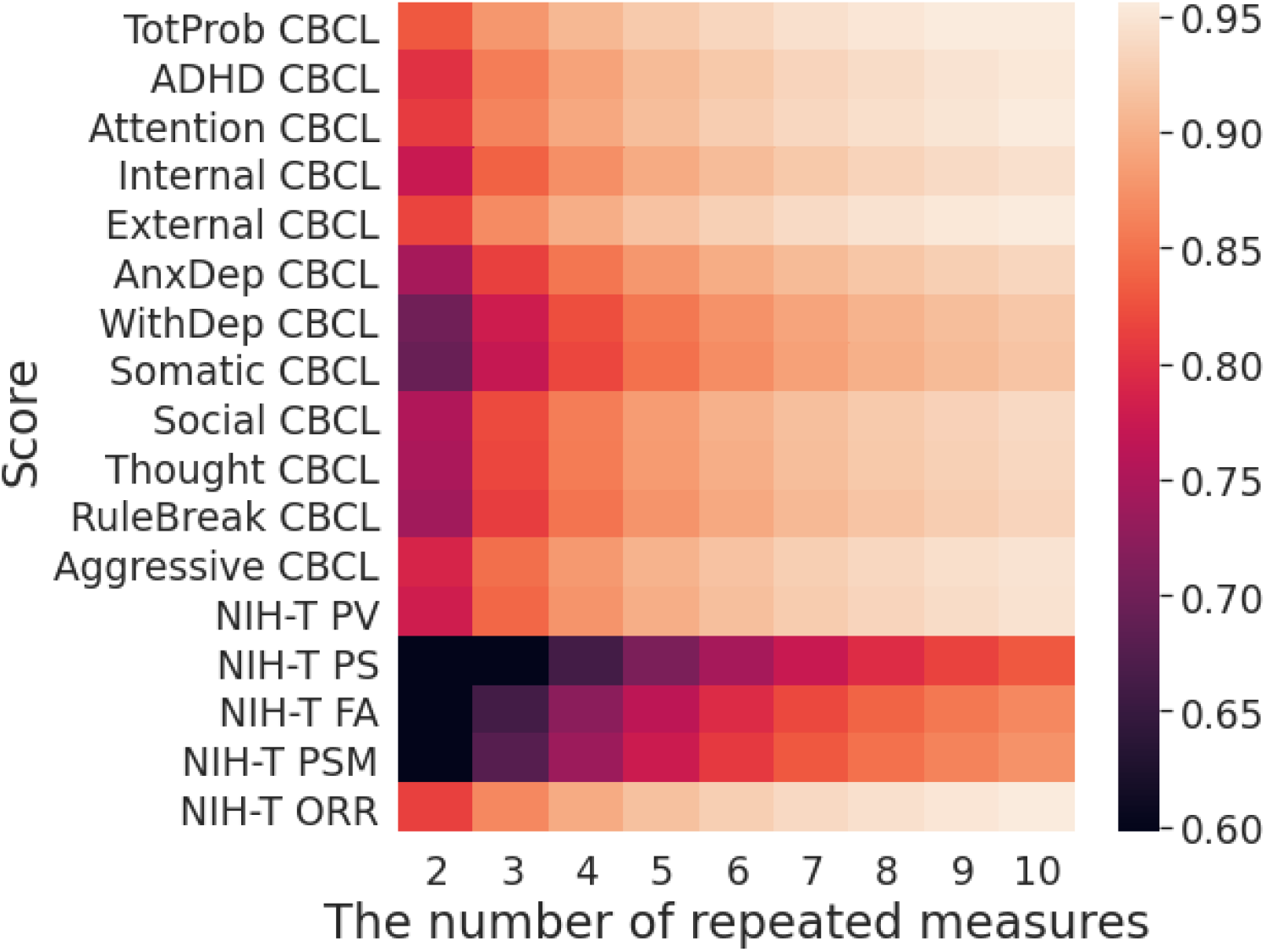
We measure the test-retest reliability for the above cognitive and clinical measures for all subjects from the ABCD study with complete data for years one and two (n = 7,249). This heatmap shows the estimated improvements in reliability that can be expected by using repeated measurements. As the number of measurements increases, all measures show large improvements in reliability. NIH-T: NIH Toolbox. PV: Picture Vocabulary Test. PS: Pattern Comparison Processing Speed Test. FA: Flanker Inhibitory Control and Attention Test. PSM: Picture Sequence Memory Test. ORR: Oral Reading Recognition Test.

**S. Figure 7.**
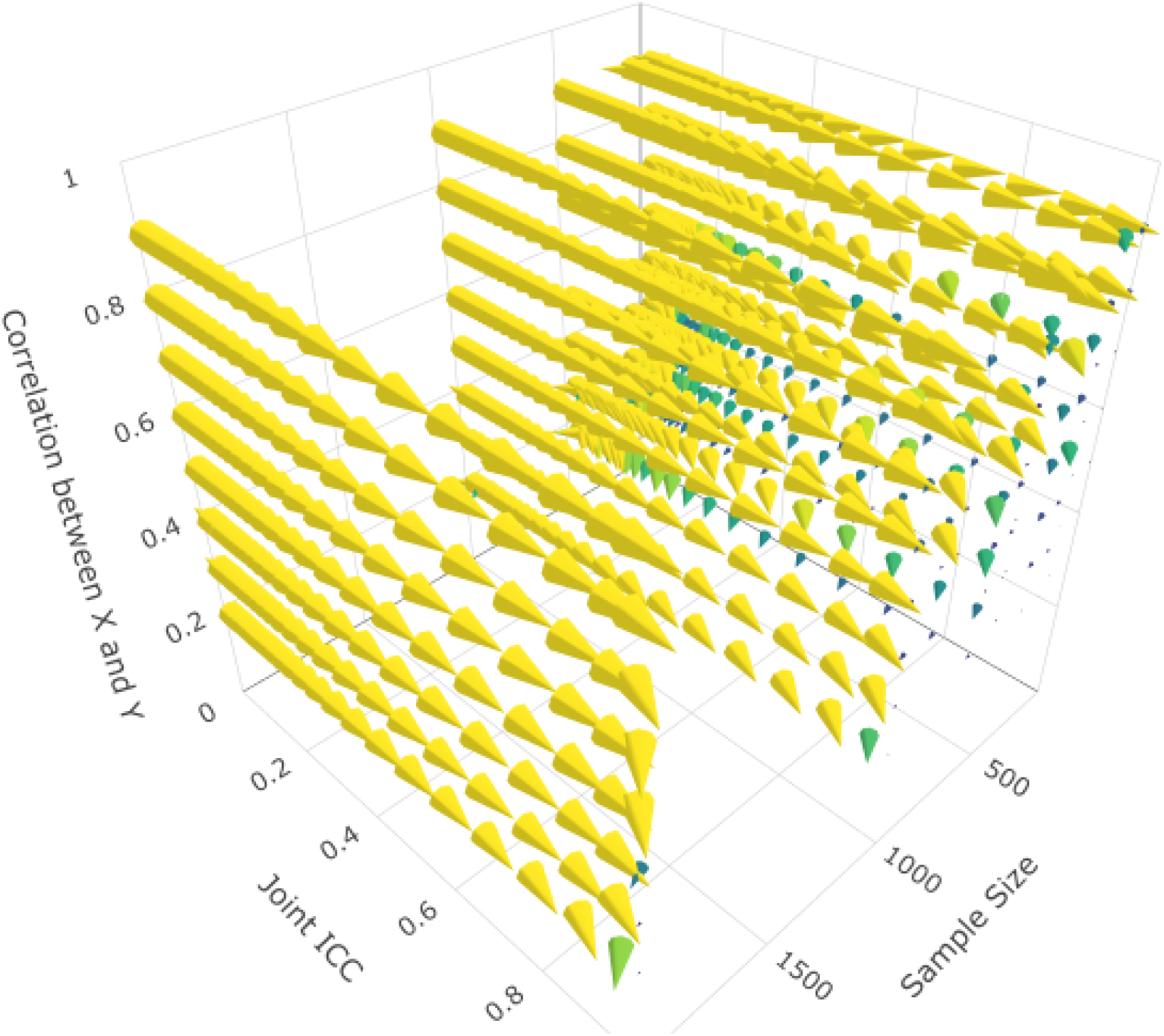
This point vector cloud informs researchers whether increasing either sample size or joint ICC (yellow) will lead to greater reductions in MSE. The color spectrum and orientation of the cones correspond to whether larger decreases in MSE are achieved by increasing sample size (blue), or increasing joint ICC (yellow). The Z axis shows how this changes as a function of the strength of relationship between X & Y. Size of cones indicate only the strength of the advantage of increasing either sample size by one level (i.e., from 200 to 500 subjects) or joint reliability by 0.1. At low sample sizes (n = 100), increasing sample size (i.e., from 100 to 200 subjects will lead to larger decreases in MSE than increasing in joint reliability. Stronger effects always benefit more from improving reliability over increasing sample size. In general, as sample size increases, increasing reliability matters more than increasing sample sizes regardless of effect size and joint ICC level. Sample sizes simulated here: n = 100, 200, 500, 1,000, 2,000.

**S. Figure 8.**
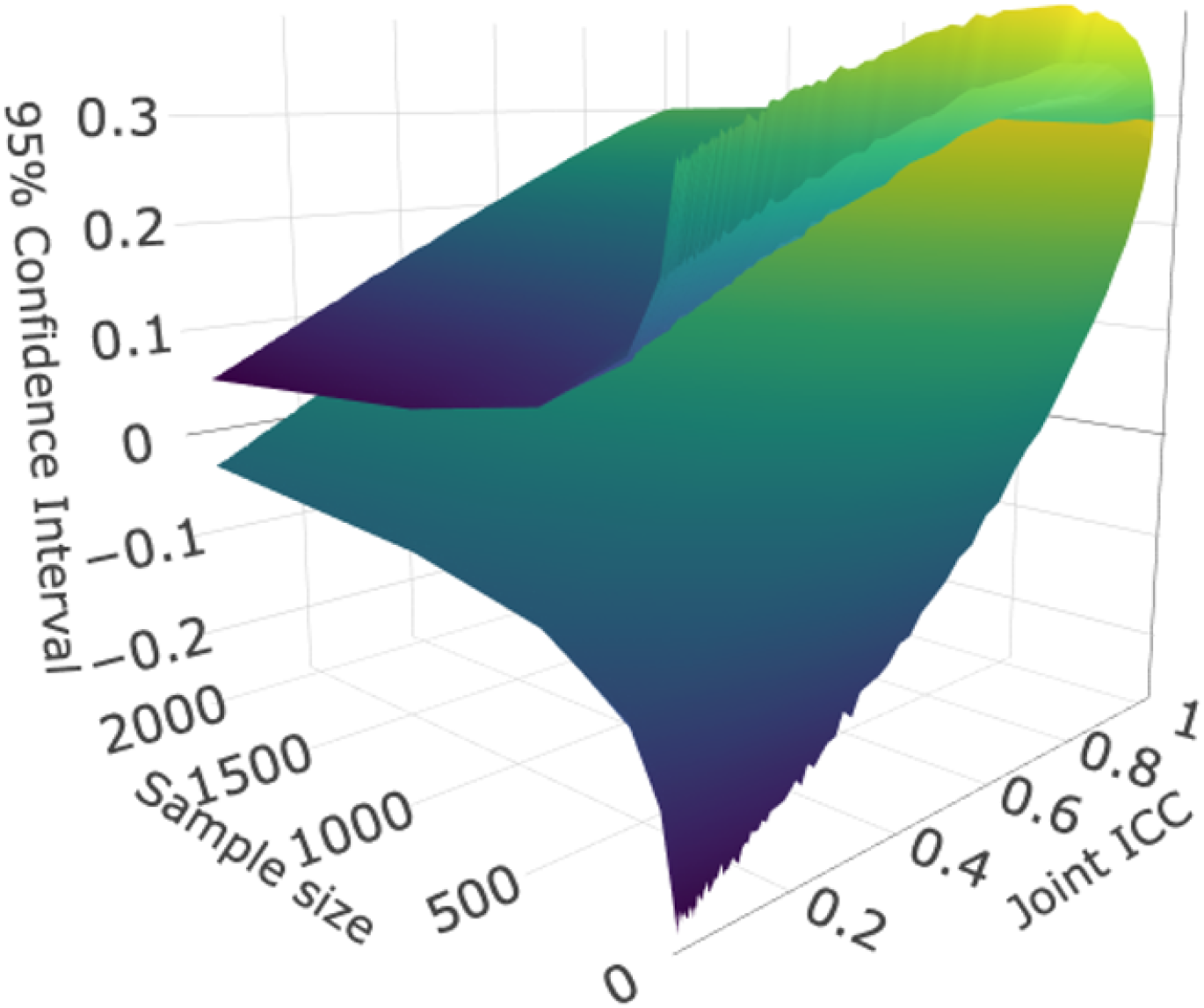
The 95% confidence interval in estimated effect sizes as a function of sample size and joint ICC. As joint ICC increases, effect size attenuation decreases and the upper and lower bounds converge on the true effect size (r = 0.3). Increasing sample sizes decreases the range of the upper and lower bounds of estimated effect sizes, with most of the decrease in variability coming from increasing samples from 0 to 500 subjects. This plot shows that increasing sample sizes when joint reliability is low (<0.2) can still produce effect sizes that are not reproducible (i.e., lower bounds that cross zero).

